# Effect of dietary Chitosan supplementation on Intestinal Barrier Function and Growth Performance in weaned piglets challenged by *Enterohemorrhagic haemolytic Escherichia coli*

**DOI:** 10.64898/2026.03.23.713631

**Authors:** Jiahao Liu, Simona De Blasio, Kunhong Xie, Xiang Li, Yuheng Luo, Ping Zheng, Xiangbing Mao, Hui Yan, Quyuan Wang, Liam Good, Ludovic Pelligand, Jun He

## Abstract

*Enterohemorrhagic Escherichia coli* O157:H7 (EHEC) is an important zoonotic pathogen that disrupts intestinal epithelial barrier integrity and induces excessive inflammatory responses, thereby leading to impaired growth performance and intestinal injury. EHEC is also an important cause Hemolytic Uremic Syndrome (HUS) in children and older adults. In pig production, chitosan is considered a promising alternative to antibiotics due to its bioadhesive and antimicrobial properties, but the effects and underlying mechanisms of chitosan (COS) under pathogenic challenge remain to be elucidated. One hundred and eight pigs were randomly divided into three treatments: an unchallenged control group (CON), an EHEC-challenged control group (ECON), and an EHEC-challenged group supplemented with 100 mg/kg COS (ECOS). Results show that EHEC challenge increased the feed conversion ratio (FCR), increased inflammatory cytokine levels, disrupted intestinal morphology, and downregulated tight junction and nutrient transporter gene expression (*P*<0.05). Dietary COS supplementation significantly improved average daily gain (ADG) and FCR during day 6-14 (*P*<0.05). Moreover, COS reduced fecal shedding of total E. coli (*P* = 0.085) and EHEC, attenuated systemic inflammation by decreasing serum TNF-α and IL-6 levels, and enhanced humoral immunity as indicated by increased IgA and IgM concentrations (*P*<0.05). Importantly, COS alleviated EHEC-induced intestinal injury by restoring villus height and villus-to-crypt ratio, with enhanced mucosal digestive enzyme activities, and upregulated expression of tight junction proteins (*ZO-1* and *occludin*) and nutrient transporters (*SGLT-1* and *PEPT1*) (*P*<0.05). In conclusion, these findings indicate that dietary COS improves growth performance in EHEC-challenged weaned pigs, with enhanced intestinal barrier integrity and nutrient transport capacity.

## Background

Weaning stress represents a critical bottleneck in early post-weaning growth and development of piglets and is often accompanied by impaired intestinal barrier function and immunosuppression, thereby markedly increasing susceptibility to enteric pathogen infection. *Enterohemorrhagic Escherichia coli* O157:H7 (EHEC), an important zoonotic pathogen, can specifically adhere to the intestinal epithelial surface and secrete virulence factors, including Shiga toxins and intimin, which disrupt epithelial structural integrity. In addition, EHEC infection activates the NLRP3 signaling pathway, triggering excessive inflammatory responses characterized by elevated production of pro-inflammatory cytokines such as TNF-α and IL-6, ultimately leading to diarrhea, intestinal injury, and even mortality in piglets [1–3]. Moreover, EHEC exhibits strong environmental survival and prolonged fecal shedding, facilitating transmission within swine herds and dissemination through the food chain, thereby posing dual threats to swine production and public health [4]. Therefore, effectively reducing intestinal colonization and shedding of EHEC at the source of pig production is of great importance for safeguarding animal and public health.

Functional polysaccharide feed additives attract increasing attention as promising alternatives to antibiotics for improving intestinal health in animal nutrition, owing to their favorable biosafety, biodegradability, and multi-target regulatory potential [5, 6]. Chitosan, is a natural cationic polysaccharide, isloated as a biproduct from crustacea or fungi, is widely available and exhibits diverse biological activities, including antibacterial properties, adsorption of harmful substances, and improvement of intestinal function [7]. Current research has predominantly examined low molecular weight chitosan (LMWC) and chitosan oligosaccharides; however, their rapid diffusion or absorption (short-chain oligosaccharides) in the intestinal lumen may restrict their capacity to directly protect the intestinal barrier under pathogenic infection [8, 9, 13] and may pose systemic exposure risks. In contrast, higher molecular weight chitosan possesses a higher overall positive charge and pronounced bioadhesive properties, enabling the formation of a stable local functional layer on microbial and or intestinal mucosal surfaces [10, 11, 12]. Under the combined challenges of weaning stress and EHEC infection, higher molecular weight chitosan may exert more targeted or effective intestinal protective effects by interfering with pathogen adhesion and colonization, enhancing intestinal barrier integrity, and modulating inflammatory responses [14, 15]. Nevertheless, systematic studies evaluating the effects of higher molecular weight chitosan (250-500 kDa) on growth performance and intestinal health in weaned piglets challenged with EHEC, as well as the underlying mechanisms, remain limited. Therefore, the present study was designed to address these gaps and to clarify the application potential and mechanisms of action of HMWC, providing a scientific basis for its use in preventing EHEC infection and maintaining intestinal health in swine production.

## Materials and methods

### Bacterial strain and culture

EHEC serotype O157:H7 (CICC 10907) was acquired from the China Institute of Veterinary Drug Control (Beijing, China). Luria-Bertani (LB) broth and LB agar were prepared and autoclaved at 121 °C, 0.11 MPa for 20 min (pH 6). The strain was revived in 10 mL LB broth at 37 °C with shaking for 24 h, then streaked onto LB agar. A single colony was inoculated into 50 mL LB broth, cultured overnight ( 37 °C and 250 rpm), followed by subculture and serial dilution on LB agar for bacterial enumeration.

### Experimental design and feed

A total of 108 piglets (Duroc⊆Landrace⊆Yorkshire) weaned at 30 days of age were randomly allocated to three treatments (18 replicates per treatment were used, and each replicate consists of 2 piglets, for each treatment the male:female ratio was 1:1): CON (pigs were fed with a basal diet), ECON (pigs were fed with a basal diet and challenged by EHEC), ECOS (pigs were fed with a basal diet containing 0.1% Chitosan (COS) product and challenged by EHEC). The COS was purchased as food grade and GMP certified. It was a white to light yellow, odourless free-flowing powder, 90.34% degree of deacetylation, with a molecular weight between 250-500 kDa. These piglets were followed for 21 days post-weaning. On day 6 post-weaning, the challenged groups received an oral dose of 100 mL LB broth containing 1×10¹⁰ CFU/mL EHEC via orogastric gavage, while the non-challenged group was administered a same volume of sterile LB medium. Experimental weaner diets, based on a corn-soybean meal formulation, were formulated to meet the ntrient specifications outlined by the National Research Council (NRC, 2012), contained a medium-high crude protein level without antimicrobial growth promoters or feed additives, and were prepared as mash at the SICAU research farm (ingredients and calculated composition shown in supplementary table 1. COS is homogenized via a two-step process: dilution with extruded corn to form a premix, followed by mixing with other ingredients.

### Animal housing and management

Piglets were housed two per 3.6 m² in slatted pens equipped with rubber insulation boards, a three-place feeder, and two nipple drinkers, in an environmentally controlled room with mechanical ventilation; temperature (28 °C in weeks 1-2, then reduced by 1.5 °C per week to 25 °C) and relative humidity (60 ± 5%) were thermostatically controlled and monitored daily, aerial ammonia was measured monthly, and empty pens and plastic boards were used to prevent cross-contamination between groups.

### Body weight, feed intake, and feed conversion ratio

Individual pig body weight was recorded at weaning, day 7, 14, and 21 post-weaning. Average daily gain (ADG, g/pig/d) and feed conversion ratio (FCR) were calculated on a pen level. Feed allowance and refusals were recorded per pen. Average Daily feed intake (ADFI, g/pig/d) was calculated for each experimental phase.

### Faecal sampling

Faeces were sampled on pen level at day 0, 7, 14, and 21 post-weaning (challenged on day 6). The sample was put in sterile cups and kept on ice until the lab. The pooled sample was homogenized with a sterile tool and 2×1 gram of faeces were stored in 2 mL cryotubes at –20°C until analysis (e.g. *E. coli* shedding). For cultivation, only faecal samples collected at d0, d7, d14, and d21 were utilized. Measurements of *E. coli* shedding (on pen level, n = 9) were performed meeting to the method as described by Busser [16], the method consists in the isolation of haemolytic *E. coli* and enumeration of total *E. coli* (log colony forming units (CFU) per mL); the specific media required for the cultivation of *E. coli* and EHEC were procured from Haber Biotechnology Ltd (Qingdao, China). Specifically, total *E. coli* were cultured on MacConkey agar (Cat no. HB6238), while EHEC were cultured on a modified sorbitol MacConkey agar (Cat no. HB0121-3) supplemented with modified sorbitol MacConkey agar additive (Cat no. HB0121-2).

### Blood sampling

At the experiment’s end, blood specimens were harvested after 12 h fasting by jugular vein puncture (in the morning on d 21 post-weaning). Whole blood was transferred into two 10 mL vacuum collection tubes; serum was subsequently isolated by centrifugation at 3,500×g for 15 min, and the resulting serum aliquots were stored at –20 °C until subsequent biochemical analysis. Enzyme-linked immunosorbent assay (ELISA) kits were used for the quantification of serum inflammatory cytokines and immune globulins: Porcine TNF-α ELISA Kit (MM-0422O1), Porcine IL-1β ELISA Kit (MM-0418O1), Porcine IL-6 ELISA Kit (MM-0418O1), and Porcine IgA, IgG, and IgM ELISA Kits (catalog nos. MM-0905O1, MM-0403O1, MM-0402O1) were all purchased from Jiangsu Enzyme-linked Biotechnology Co., Ltd. (Jiangsu, China). The indicators were measured according to the manufacturer’s instructions. Others serum samples were transported to Ya’an People’s Hospital (Ya’an, China) for hematological parameter assessment (*e.g.* total protein, albumin, globulin, glucose).

### Tissue sampling

Intestinal tissue collection was conducted subsequent to blood sampling. Piglets were euthanized by intravenous injection of sodium pentobarbital at 200 mg/kg BW, and then the each segment of the small intestine were dissected out. The middle segments of the intestinal tissues 4 cm obtained were gently flushed with ice-cold PBS, followed by fixation in 4% paraformaldehyde solution for morphological analyses. The intestinal mucosa was obtained from the residual intestinal segments with scalpel blade and place in frozen tube then frozen by immersion in liquid nitrogen and stored at –80 °C until analysis (e.g. lactase, amylase, sucrase, alkaline phosphatase, endotoxin, and nutrient absorption and transportation related expression gene expression levels).

**(1)** **Intestinal morphology:** Fixed small intestinal segments were dewaxed with graded anhydrous ethanol, stained by hematoxylin and eosin (H&E), and sealed with neutral resin as described previously [17]. The sections were then analyzed for crypt depth (CD), villus height (VH), and VH/CD ratio using an Image-Pro Plus 6.0 image analysis system (Image-Pro Plus 6.0, Media Cybernetics, Maryland, USA.
**(2)** **Enzyme activity and endotoxin:** The intestinal mucosa was homogenized with saline, and then the supernatants were isolated and utilized to determine the enzyme activities such as the lactase (Porcine lactase ELISA Kit MM-MM-1746O1), invertase (Porcine invertase ELISA Kit MM-1747O1), and maltase (Porcine maltase ELISA Kit MM-1748O1), Alkaline phosphatase activity (Porcine ALP ELISA Kit MM-36368O1) and endotoxin content (Porcine ET ELISA Kit MM-36368O1) were also determined by using specific enzyme-linked immunosorbent assay kits (Jiangsu Meimian industrial Co., Ltd.). All procedures were performed according to the instructions.
**(3)** **Q-PCR quantification:** Total mucosal RNA was extracted with TRIzol and reverse-transcribed to cDNA using a PrimeScript™ RT kit with gDNA Eraser (Takara), under conditions of 37 °C for 15 min and 85 °C for 5 s. qPCR was performed with SYBR® Green reagents (Takara) using the program: 95 °C for 30 s, 40 cycles of 95 °C for 5 s and 60 °C for 30 s. Target gene expression was analyzed by the 2^-ΔΔCt^ method with β-actin as the internal control [18], and all primers are listed in supplementary table 2.

### Statistical analysis

The experimental data was analysed using one-way analysis of variance (ANOVA) via the General Linear Model function of SPSS 24.0 (SPSS, Inc). The analyses were performed with and without excluding outliers. Any outliers removed prior to statistical analyses were indicated. Significant differences were declared at *P*<0.05, with near significant trends as 0.05<*P*≤0.10. (Near) Significant fixed effects were further analysed by Least Significant Differences (LSD, Fisher’s LSD method) to compare treatment means. Data is presented as absolute means. The *P*-value, LSD, and standard error of the mean (SEM) are reported per response parameter.

### Animal ethics

This study was conducted in strict compliance with the animal welfare regulations of the People’s Republic of China and was designed to mitigate any potential pain or distress inflicted upon experimental animals, consistent with the research’s scientific aims. The experimental protocol was approved by the Institutional Animal Care and Use Committee of Sichuan Agricultural University. The use of COS was approved by the China Ministry of Agriculture. The study was also reviewed and approved by RVC’s Animal Welfare and Ethical Review Body (AWERB URN 2022-0166N).

### Unexpected issues

There were no unexpected issues excepting the African Swine Fever. However, we have collected saliva samples from pigs and sent it for virus detection (samples were detected by authoritative testing organization), and only non-infected pigs were allowed to enter the research farm. Moreover, stringent disinfection procedures and management were performed throughout the trial.

## Results

### Animal health and mortality

A total of 108 piglets were included in this experiment at the weaning stage and were followed up for 3 weeks. During the experimental period, all pigs were maintained in normal feeding, drinking and mental status without any medical intervention, and no deaths occurred (zero mortality).

### Growth Performance and faecal consistency

As shown in Table 1, the FCR was significantly higher in the ECON group than in the CON and ECOS group, and the ADG was significantly lower in ECON group than in the ECOS group during day 6-14 (*P*<0.05). However, there were no differences in ADFI and BW amongst the three group during the experiment period.

**Table 1.**
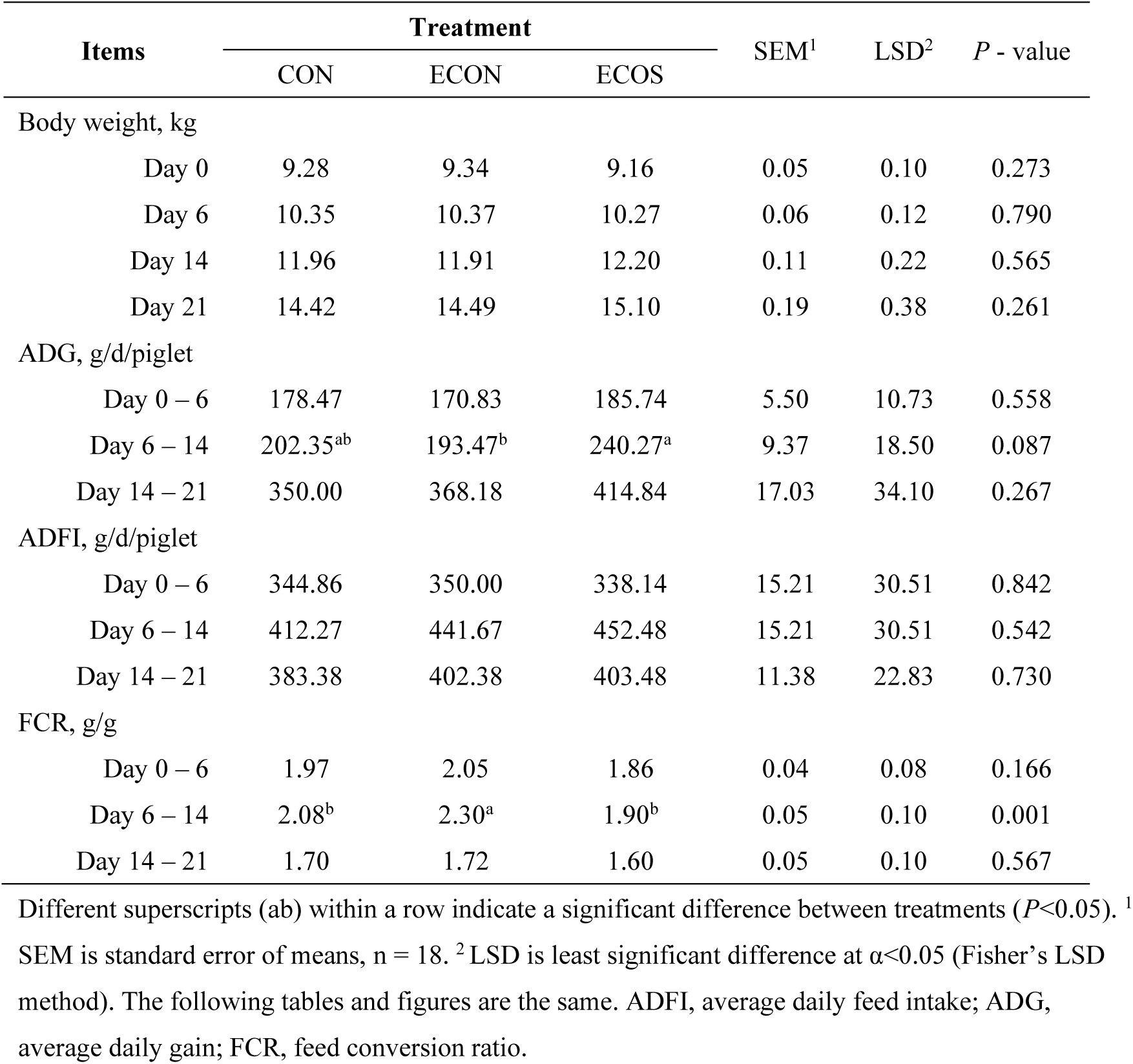
Effect of COS on growth performance and faecal consistency scores in weaned pigs upon EHEC Challenge.

### E. coli and haemolytic E. coli shedding

Pigs in the ECON and ECOS groups were challenged by EHEC at day 6 (the CON group was infused by LB culture medium). As shown in Figure 1, there were no differences in the shedding of total *E coli* and EHEC amongst the three groups at day 0. Maximal shedding in the ECON and ECOS groups was observed at day 7, and the shedding of total *E coli* and EHEC continuously decreased from day 14 to day 21. As compared to the ECON group, dietary COS supplementation decreased shedding of the total *E coli* and EHEC in the ECOS group at day 7 (*P* = 0.085) and day 14 (*P*>0.05) (Figure 1). However, there were no significant differences amongst the three group (This may attribute to the big variations during the cultivation).

**Figure 1.**
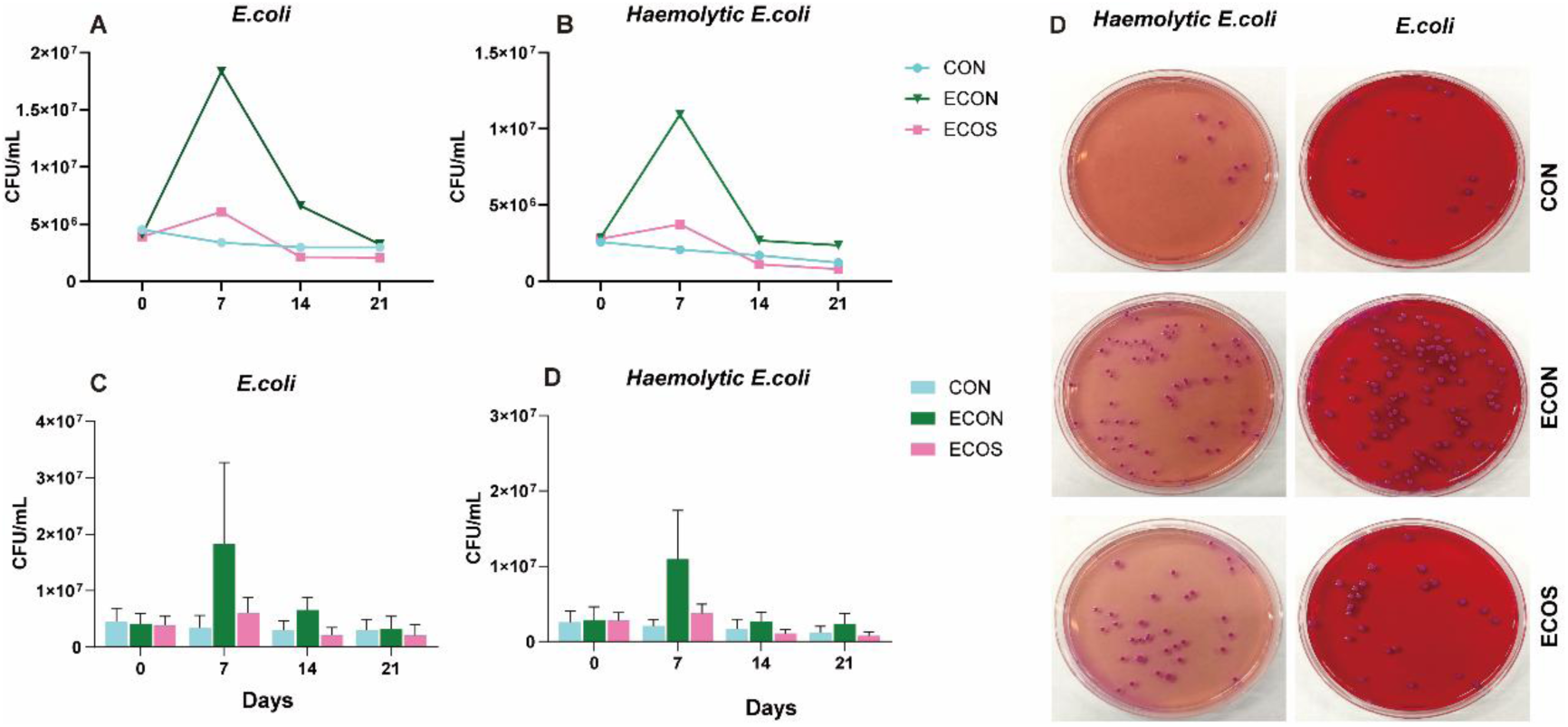
Effects of COS on CFU counts of rectal total *E. coli* and *haemolytic E. coli* shedding in weaned pigs after EHEC challenge. Panels (A), (B) and (C), (D) show the growth of these two *E. coli* populations from 0 to 21 days respectively; (E) illustrates the growth of the two populations on selective media at day 7, following 10^4^, 10^5^ and 10^6^ fold dilutions. There were no statistical differences between each treatment group.

### Clinical serum biochemical parameters

As shown in Table 2, there were no differences in the serum concentrations of serum albumin, globulin, and glucose amongst the three groups. However, EHEC challenge decreased the serum concentration of TBILI, and decreased the serum activities of AST and CK (*P* < 0.05).

**Table 2.**
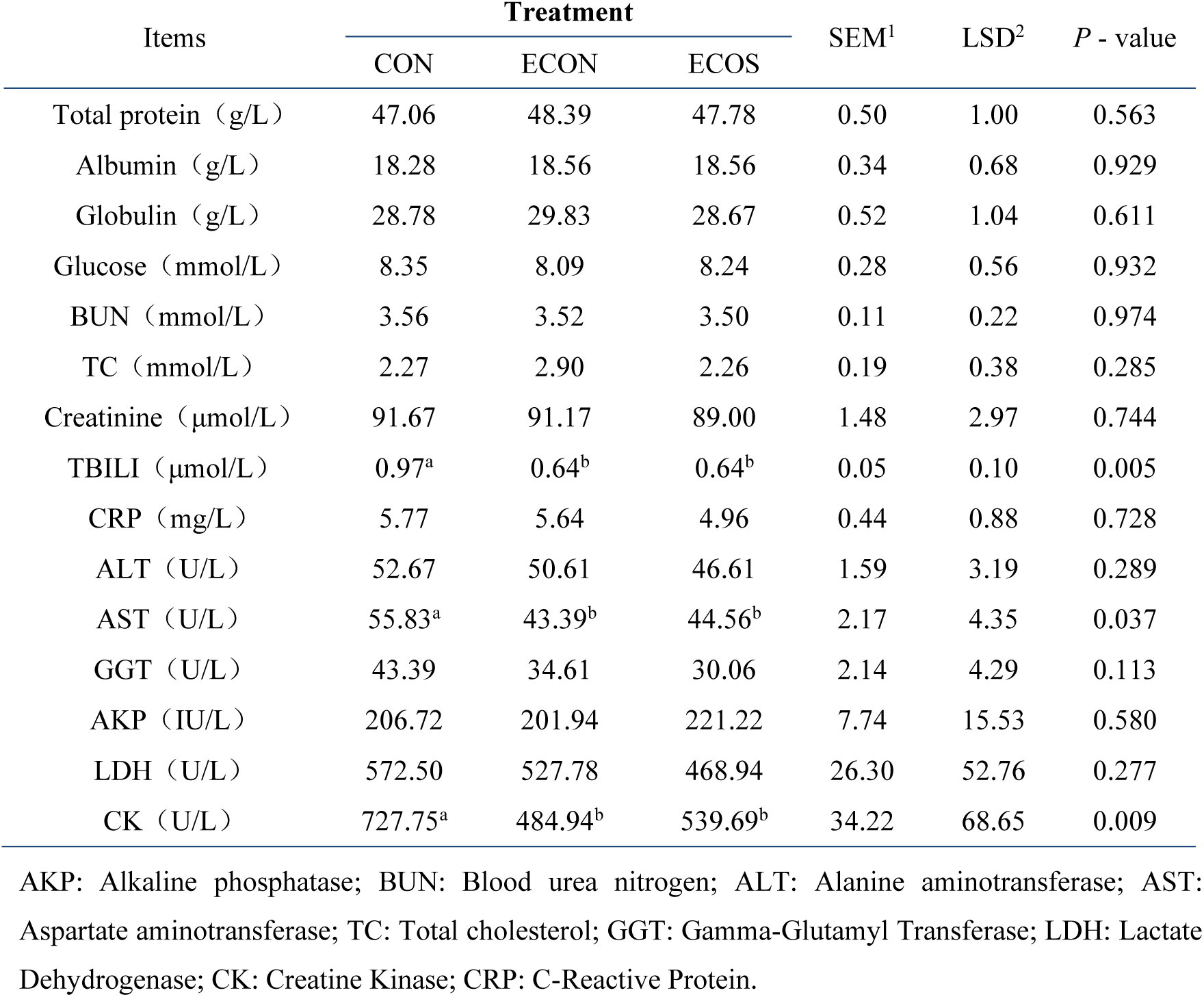
Effect of COS on clinical biochemical parameters upon EHEC challenge.

| Items | Treatment |  |  | SEM<sup>1</sup> | LSD<sup>2</sup> | *P* - value |
| --- | --- | --- | --- | --- | --- | --- |
|  | CON | ECON | ECOS |  |  |  |
| Total protein (g/L) | 47.06 | 48.39 | 47.78 | 0.50 | 1.00 | 0.563 |
| Albumin (g/L) | 18.28 | 18.56 | 18.56 | 0.34 | 0.68 | 0.929 |
| Globulin (g/L) | 28.78 | 29.83 | 28.67 | 0.52 | 1.04 | 0.611 |
| Glucose (mmol/L) | 8.35 | 8.09 | 8.24 | 0.28 | 0.56 | 0.932 |
| BUN (mmol/L) | 3.56 | 3.52 | 3.50 | 0.11 | 0.22 | 0.974 |
| TC (mmol/L) | 2.27 | 2.90 | 2.26 | 0.19 | 0.38 | 0.285 |
| Creatinine (μmol/L) | 91.67 | 91.17 | 89.00 | 1.48 | 2.97 | 0.744 |
| TBILI (μmol/L) | 0.97<sup>a</sup> | 0.64<sup>b</sup> | 0.64<sup>b</sup> | 0.05 | 0.10 | 0.005 |
| CRP (mg/L) | 5.77 | 5.64 | 4.96 | 0.44 | 0.88 | 0.728 |
| ALT (U/L) | 52.67 | 50.61 | 46.61 | 1.59 | 3.19 | 0.289 |
| AST (U/L) | 55.83<sup>a</sup> | 43.39<sup>b</sup> | 44.56<sup>b</sup> | 2.17 | 4.35 | 0.037 |
| GGT (U/L) | 43.39 | 34.61 | 30.06 | 2.14 | 4.29 | 0.113 |
| AKP (IU/L) | 206.72 | 201.94 | 221.22 | 7.74 | 15.53 | 0.580 |
| LDH (U/L) | 572.50 | 527.78 | 468.94 | 26.30 | 52.76 | 0.277 |
| CK (U/L) | 727.75<sup>a</sup> | 484.94<sup>b</sup> | 539.69<sup>b</sup> | 34.22 | 68.65 | 0.009 |

AKP: Alkaline phosphatase; BUN: Blood urea nitrogen; ALT: Alanine aminotransferase; AST: Aspartate aminotransferase; TC: Total cholesterol; GGT: Gamma-Glutamyl Transferase; LDH: Lactate Dehydrogenase; CK: Creatine Kinase; CRP: C-Reactive Protein.

### Serum inflammatory cytokines,immunoglobulins, and endotoxins

As shown in Table 3, EHEC challenge significantly increased the serum concentration of TNF-α. However, dietary COS supplementation not only decreased the serum concentrations of TNF-α and IL-6, but also increased concentrations of immunoglobulins such as the IgA and IgM in the EHEC-challenged pigs (*P*<0.05).

**Table 3.** Effect of COS on serum inflammatory cytokines, immunoglobulins, and endotoxin in weaned pigs upon EHEC challenge.

| Items | Treatment |  |  | SEM<sup>1</sup> | LSD<sup>2</sup> | *P* - value |
| --- | --- | --- | --- | --- | --- | --- |
|  | CON | ECON | ECOS |  |  |  |
| TNF-α (pg/ml) | 47.91<sup>b</sup> | 75.16<sup>a</sup> | 43.27<sup>b</sup> | 4.40 | 8.83 | 0.001 |
| IL-1β (ng/L) | 65.09 | 73.00 | 65.39 | 3.60 | 7.22 | 0.604 |
| IL-6 (ng/L) | 263.3<sup>ab</sup> | 290.93<sup>a</sup> | 231.09<sup>b</sup> | 8.92 | 17.89 | 0.020 |
| IgG (μg/mL) | 141.80 | 133.74 | 102.39 | 9.38 | 18.82 | 0.196 |
| IgA (μg/mL) | 13.47<sup>a</sup> | 9.67<sup>b</sup> | 13.88<sup>a</sup> | 0.71 | 1.42 | 0.025 |
| IgM (μg/mL) | 11.96<sup>ab</sup> | 9.95<sup>b</sup> | 14.83<sup>a</sup> | 0.82 | 1.64 | 0.047 |

### Intestinal morphology

As shown in Table 4 and Figure 2, EHEC challenge decreased the V/C in small intestine (duodenum, jejunum, and ileum). However, dietary COS supplementation significantly increased villus height and the V/C in the ECOS group (*P* < 0.05).

**Figure 2.**
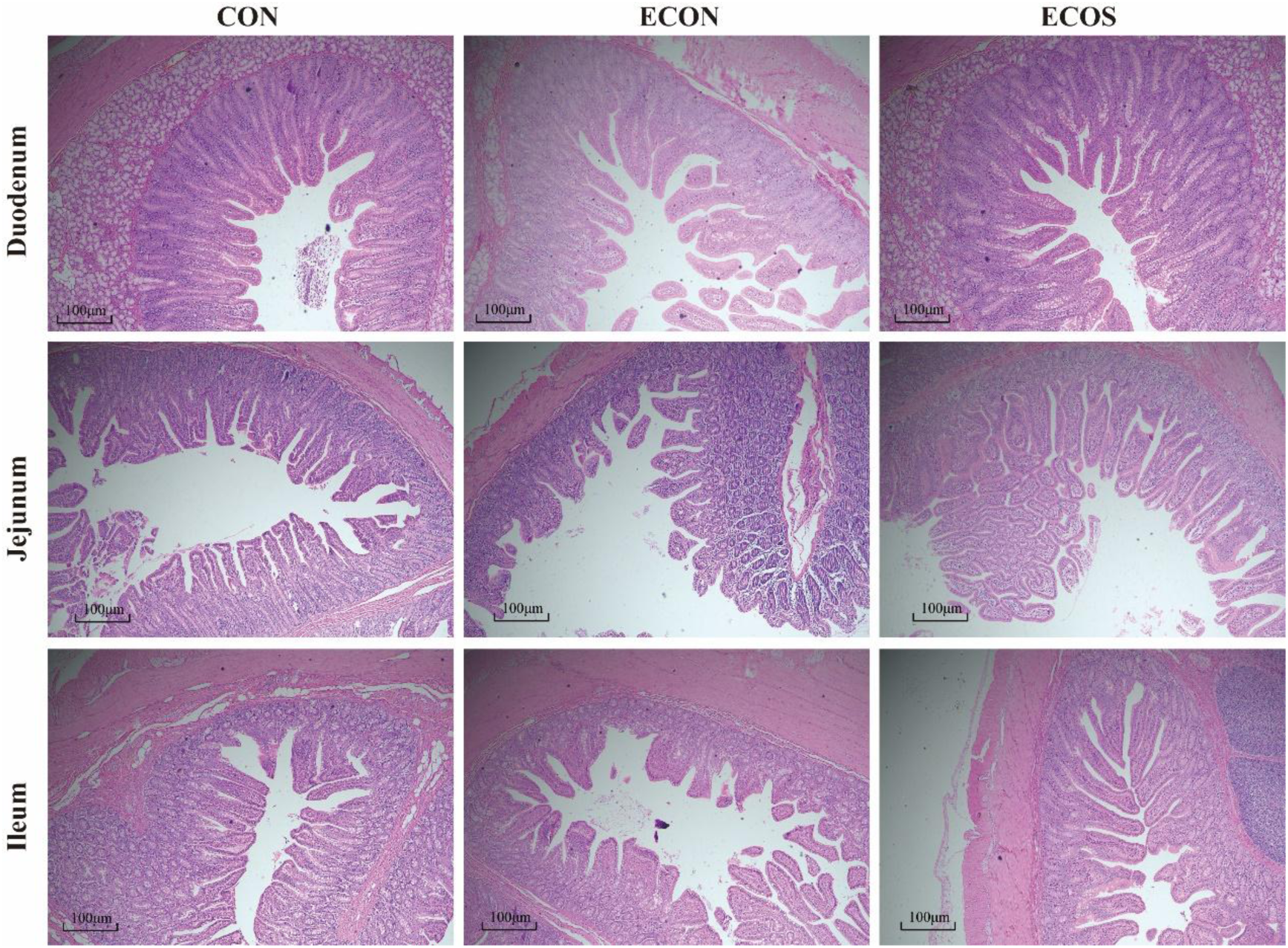
Representative exmaples of intestinal morphology in weaned pigs upon EHEC challenge (H&E × 40, represent pictures for the villus). Duodenum, jejunum and ileum in the CON group revealed a normal appearance with regular intestinal villus structure. However, in the ECON group, the intestinal lesions were obvious and some intestinal villi were necrotic and shed, or even disappeared, especially in the jejunum. In the duodenum, jejunum and ileum in the ECOS group there were no obvious intestinal lesions and only a few of the intestinal villi were necrotic and shed.

**Table 4.** Effect of COS on intestinal morphology in weaned pigs upon EHEC challenge.

| Items | Treatment |  |  | SEM<sup>1</sup> | LSD<sup>2</sup> | *P* - value |
| --- | --- | --- | --- | --- | --- | --- |
|  | CON | ECON | ECOS |  |  |  |
| **Duodenum** |  |  |  |  |  |  |
| Villus height (μm) | 553.97<sup>a</sup> | 513.41<sup>b</sup> | 597.64<sup>a</sup> | 10.01 | 20.08 | 0.001 |
| Crypt depth (μm) | 365.92<sup>ab</sup> | 413.85<sup>a</sup> | 338.00<sup>b</sup> | 9.26 | 18.58 | 0.021 |
| V/C | 1.64<sup>a</sup> | 1.33<sup>b</sup> | 1.84<sup>a</sup> | 0.04 | 0.08 | 0.001 |
| **Jejunum** |  |  |  |  |  |  |
| Villus height (μm) | 491.34<sup>a</sup> | 434.46<sup>b</sup> | 517.64<sup>a</sup> | 9.24 | 18.54 | 0.001 |
| Crypt depth (μm) | 239.62<sup>b</sup> | 323.21<sup>a</sup> | 252.75<sup>b</sup> | 7.42 | 14.88 | 0.001 |
| V/C | 2.19<sup>a</sup> | 1.40<sup>b</sup> | 2.19<sup>a</sup> | 0.06 | 0.12 | 0.001 |
| **Ileum** |  |  |  |  |  |  |
| Villus height (μm) | 482.52<sup>a</sup> | 350.00<sup>b</sup> | 440.55<sup>a</sup> | 10.87 | 21.81 | 0.001 |
| Crypt depth (μm) | 207.84<sup>b</sup> | 226.86<sup>a</sup> | 192.36<sup>b</sup> | 3.96 | 7.94 | 0.001 |
| V/C | 2.33<sup>a</sup> | 1.62<sup>b</sup> | 2.35<sup>a</sup> | 0.07 | 0.14 | 0.001 |

V/C: the ratio of villus height to crypt depth.

### Expressions of epithelial cytoskeleton-related genes

As shown in Figure 3, EHEC challenge significantly downregulated the expression levels of *itga5* and *itgb1* in the jejunal mucosa (*P*<0.05). Dietary COS supplementation alleviated these changes by significantly increasing *itga5* and *itgb1* expression in ECOS group (*P*<0.05).

**Figure 3.**
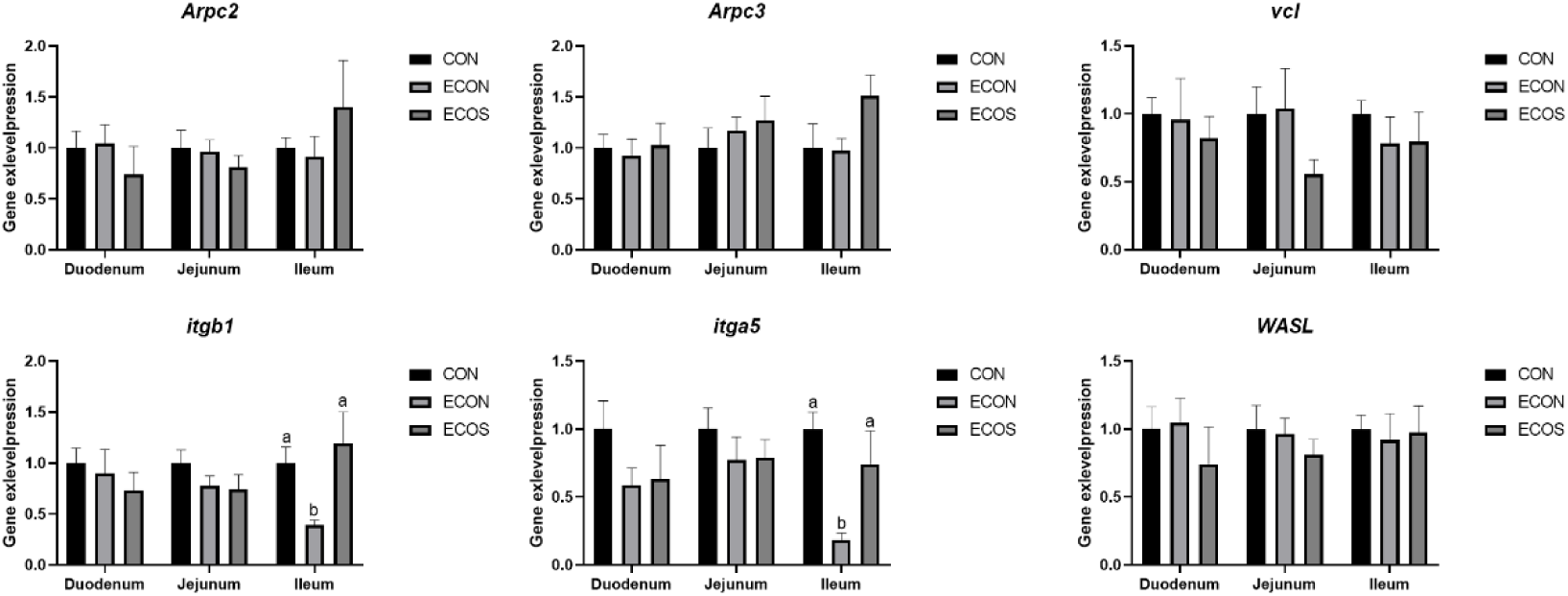
Effects of COS on the expression of cytoskeleton-related genes genes; itga5: integrin alpha-5; itgb1: integrin beta-1; vcl: vinculin; WASL: wiskott–aldrich syndrome protein-like (n-wasp); Arpc2: actin-related protein 2/3 complex subunit 2; Arpc3: actin-related protein 2/3 complex subunit 3.

### Intestinal mucosal enzyme activity

As shown in Table 5, EHEC challenge resulted in decreases in the activities of alkaline phosphatase (ALP), maltase, invertase, and lactase in the jejunal mucosa. However, dietary COS supplementation significantly increased their activities in the ECOS group (*P*<0.05).

**Table 5.** Effect of COS on intestinal mucosal enzyme activity upon EHEC challenge.

| Items | Treatment |  |  | SEM<sup>1</sup> | LSD<sup>2</sup> | *P* - value |
| --- | --- | --- | --- | --- | --- | --- |
|  | CON | ECON | ECOS |  |  |  |
| **Duodenum** |  |  |  |  |  |  |
| ALP (ng/L) | 67.58 | 77.94 | 71.25 | 2.78 | 5.58 | 0.310 |
| Maltase (ng/L) | 495.79 | 433.33 | 429.91 | 26.90 | 53.96 | 0.540 |
| Invertase (ng/L) | 212.10 | 212.05 | 259.84 | 13.11 | 26.30 | 0.229 |
| Lactase (pg/ml) | 179.69 | 188.05 | 193.51 | 9.92 | 19.90 | 0.854 |
| ET (ng/L) | 109.08 | 116.21 | 113.23 | 5.41 | 10.85 | 0.868 |
| **Jejunum** |  |  |  |  |  |  |
| ALP (ng/L) | 102.33<sup>a</sup> | 64.28<sup>b</sup> | 77.27<sup>b</sup> | 3.79 | 5.58 | 0.001 |
| Maltase (ng/L) | 366.09<sup>a</sup> | 289.59<sup>b</sup> | 353.28<sup>a</sup> | 12.77 | 53.96 | 0.029 |
| Invertase (ng/L) | 256.62<sup>a</sup> | 176.66<sup>b</sup> | 226.82<sup>a</sup> | 8.43 | 31.69 | 0.001 |
| Lactase (pg/ml) | 196.92<sup>a</sup> | 136.73<sup>b</sup> | 174.23<sup>a</sup> | 7.64 | 19.90 | 0.003 |
| ET (ng/L) | 127.68 | 108.66 | 109.61 | 4.61 | 10.85 | 0.297 |
| **Ileum** |  |  |  |  |  |  |
| ALP (ng/L) | 32.48 | 29.98 | 32.75 | 2.08 | 4.17 | 0.840 |
| Maltase (ng/L) | 286.48 | 310.67 | 286.73 | 11.76 | 23.59 | 0.637 |
| Invertase (ng/L) | 147.02 | 150.37 | 162.61 | 3.98 | 7.98 | 0.246 |
| Lactase (pg/ml) | 197.01 | 210.03 | 230.25 | 3.98 | 7.98 | 0.298 |
| ET (ng/L) | 101.03 | 105.04 | 117.68 | 2.80 | 5.62 | 0.128 |

ALP: Alkaline Phosphatase; ET: Endotoxin.

### Expressions of crtical functional genes

As shown in Figure 4, EHEC challenge downregulated the nutrient transporters and intestinal barrier related genes expression levels such as the *SGLT-1*, *ZO-1* and *Occludin* in the jejunal mucosa (*P*<0.05). However, dietary COS supplementation significantly upregulated their expression levels in the ECOS group (*P*<0.05). Dietary COS supplementation elevated the jejunal *GLUT-2* and *PETP1* gene expression levels (*P*<0.05).

**Figure 4.**
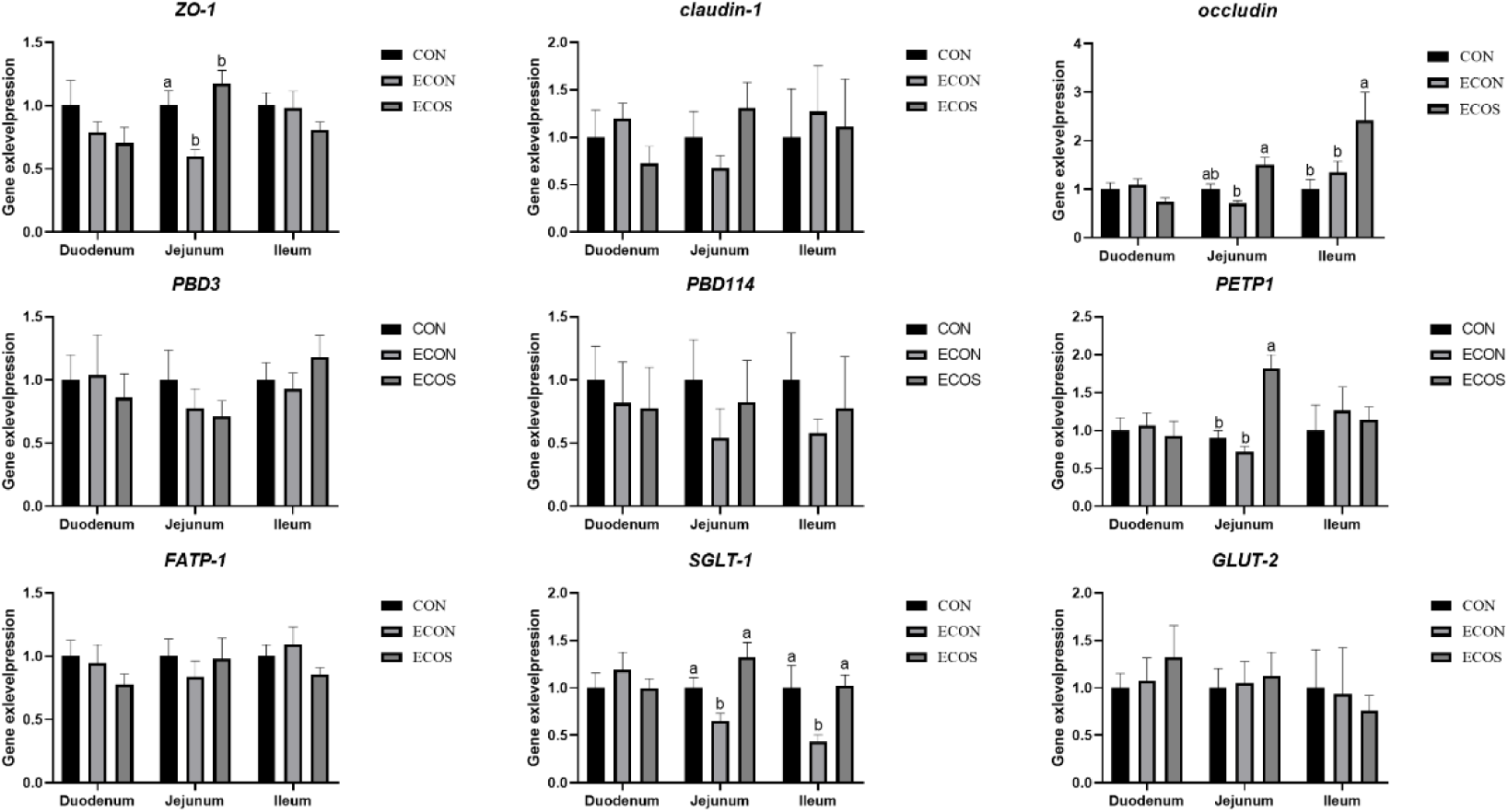
Effects of COS on the expression of intestinal epithelial barrier-related genes; *GLUT-2*: Glucose Transporter-2; *SGLT-1*: Sodium-Glucose Linked Transporter-1; *FATP-1*: Fatty Acid Transport Protein-1; *ZO-1*: Zonula Occludens-1.

## Discussion

Since its first identification during a foodborne disease outbreak in the United States in 1982, EHEC has been widely recognized as an important zoonotic enteric pathogen, capable of causing severe diarrhea and hemorrhagic colitis and posing a persistent threat to livestock production and public health risks, particularly to children and older adults [19]. Weaned piglets, whose intestinal structure and function are not yet fully developed and whose mucosal barrier and immune defenses remain immature, are highly susceptible to enteric pathogens such as EHEC and therefore represent an appropriate model for investigating strategies aimed at limiting pathogen colonization. Chitosan is a natural cationic polysaccharide that is poorly digested and absorbed by animals and is characterized by a high density of positive charges and good biocompatibility. In the gastrointestinal tract, higher molecular weight chitosan is not readily degraded and so can potentially interact electrostatically with the intestinal mucosa and microbial surfaces, allowing it to establish a relatively stable local presence within the intestinal lumen that inhibits pathogen infection processes. These properties confer chitosan with unique advantages as a tool to modulate intestinal pathogen colonization and protect epithelial barrier functions. In this study, dietary COS (250-500 kDa) supplementation markedly alleviated the adverse effects of EHEC challenge on growth performance in weaned piglets. Specifically, during the post-challenge period from days 6 to 14, piglets receiving COS exhibited significantly greater ADG and a significantly lower FCR compared with the challenged group. The improvement in growth performance was accompanied by a sustained reduction in fecal shedding of total *E. coli* and hemolytic *E. coli.* Although the differences between groups did not reach statistical significance at all sampling points, a consistent and clear trend toward reduced colonization was observed in the COS –treated group on days 7 and 14 post-infection. These findings suggest that COS may reduce the detrimental effects of EHEC by interfering with pathogen colonization and persistence within the intestinal lumen. The inhibitory effects of COS on EHEC colonization are unlikely to result from a single antimicrobial mechanism but may instead involve multiple complementary modes of action. First, amino groups on the chitosan molecule chelate metal ions such as Ca²⁺ and Mg²⁺, which are essential for bacterial outer barrier integrity, cell growth and virulence expression, thereby altering the luminal microenvironment and disrupting bacterial metabolic activity and toxin production [20]. Second, as a positively charged polymer, COS can interact electrostatically with the negatively charged bacterial cell membrane, potentially promoting bacterial aggregation in the gut and reducing effective adhesion to the intestinal epithelium—an essential prerequisite for stable colonization and pathogenesis of EHEC. In addition, these interactions may compromise membrane integrity and permeability, leading to leakage of intracellular contents and impaired bacterial viability [21]. Importantly, by limiting direct contact between pathogens and the intestinal epithelium, COS may attenuate EHEC-induced mucosal inflammation and epithelial disruption, thereby reducing the energetic and nutritional costs associated with immune activation and tissue repair. Consequently, a greater proportion of nutrients can be allocated to growth-related physiological processes, ultimately resulting in improved growth performance.

Immunoglobulins are key antibacterial effector molecules of the host immune system that mediate pathogen clearance and reduce pathogenicity by forming antigen–antibody complexes and subsequently neutralizing microbial toxins, inhibiting pathogen adhesion and colonization, activating the complement system, and enhancing neutrophil– and macrophage-mediated phagocytosis [22, 23]. In the present study, COS supplementation significantly attenuated the EHEC-induced decreases in serum IgG, IgM, and IgA levels, indicating an enhanced anti-infective capacity in weaned piglets. Moreover, pathogen invasion activates inflammatory signaling pathways, leading to excessive release of pro-inflammatory cytokines to initiate immune responses, whereas sustained or excessive inflammation may exacerbate tissue damage. Our results show that EHEC infection markedly increased serum TNF-α and IL-6 levels in piglets, while dietary COS supplementation effectively suppressed the elevation of these pro-inflammatory cytokines, further suggesting a potential role of COS in maintaining immune homeostasis and alleviating inflammatory responses. These are consistent with previous studies demonstrating that enterotoxigenic Escherichia coli exacerbates inflammation via the TLR4/NF-κB signaling pathway, whereas chitosan mitigates inflammatory responses by suppressing TLR4 expression and subsequently blocking downstream inflammatory signaling cascades [24, 25].

The intestine is a key organ for nutrient digestion and absorption, and the integrity of its epithelial structure and function is needed for efficient nutrient utilization and provide a first line of defence against the invasion of harmful microorganisms [26, 27]. Previous studies have shown that ETEC can colonize the small intestine and secrete exotoxins, thereby disrupting intestinal morphology [28]. In contrast, EHEC primarily delivers effector proteins, including Tir and EspA/B/D, into host epithelial cells via the type III secretion system encoded by the locus of enterocyte effacement (LEE). These effectors mediate Intimin-Tir-dependent intimate adherence and induce actin cytoskeleton rearrangement, leading to the formation of characteristic attaching and effacing (A/E) lesions, which compromise epithelial structural integrity and exacerbate local inflammatory responses [29, 30]. In this study, EHEC challenge reduced villus height and the V/C ratio while increasing crypt depth in the small intestine of piglets, indicating a pronounced impairment of the intestinal absorptive surface area and functional status. Notably, EHEC challenge also markedly downregulated the expression of *ITGA5* and *ITGB1* in the intestinal epithelium. Integrin α5β1, encoded by *ITGA5* and *ITGB1*, is a key mediator of epithelial cell-basement membrane adhesion and plays an essential role in maintaining epithelial structural stability and villus architecture [31,32]. The decreased expression of these integrin subunits suggests that EHEC infection weakens epithelial anchorage to the basement membrane. This molecular alteration is consistent with the pathological features of A/E lesions, which involve epithelial cytoskeletal rearrangement and disruption of adhesion structures, and may, at least in part, exacerbate villus shortening and the loss of intestinal absorptive surface area. However, dietary supplementation with COS markedly alleviated these morphological alterations, as evidenced by increased villus height and V/C ratio, accompanied by a concurrent restoration of *ITGA5* and *ITGB1* expression. These findings suggest that COS may attenuate EHEC-induced A/E damage at the structural level by maintaining the stability of epithelial-basement membrane interactions. Alterations in intestinal morphology are often accompanied by changes in brush-border digestive enzyme activities. Consistent with this notion, EHEC infection significantly reduced the activities of alkaline phosphatase, maltase, sucrase, and lactase, whereas COS supplementation effectively mitigated these reductions, further supporting the role of COS in preserving overall intestinal digestive and absorptive homeostasis.

Intestinal barrier integrity and nutrient transport capacity are critical determinants of growth performance in weaned piglets under enteric pathogen challenge [33, 34]. The present results demonstrate that dietary supplementation with COS markedly preserved the expression of the tight junction proteins ZO-1 and occludin in EHEC-challenged piglets, indicating effective protection of epithelial structural integrity and a reduced risk of increased paracellular permeability and trans-epithelial passage of pathogen-associated factors. Maintenance of tight junction architecture is essential for limiting pathogen translocation and preventing excessive immune activation, processes that are widely recognized as major contributors to growth impairment during the post-weaning period [35, 36]. Moreover, COS supplementation increased the gene expression levels of the nutrient transporters *SGLT-1* and *PEPT1* in both the jejunum and colon. *SGLT-1* is the key transporter mediating the active absorption of glucose and galactose across the brush-border membrane of the small intestine, whereas PEPT1 is primarily responsible for the uptake of dipeptides and tripeptides and represents the major pathway for the absorption of protein digestion products in the intestine [37, 38]. Collectively, these findings indicate that the beneficial effects of COS are not limited to direct antimicrobial activity but also involve its function in mucosal protection within the intestinal lumen, possibly Collectively, these findings indicate that the beneficial effects of COS are not limited to direct antimicrobial activity but also involve its function in mucosal protection within the intestinal lumen, possibly by forming a layer on the mucosal surface. Higher molecular weight chitosan may provide stronger and more stable electrostatic interactions with the mucus layer and epithelial surface, helping to reduce direct contact between EHEC and epithelial cells. Such effects could in-turn stabilize tight junction structures, providing a more favourable microenvironment for the expression and function of nutrient transporters.

## Conclusion

In summary, dietary COS supplementation (100 mg/kg) improved growth performance and reduced the shedding of total *E coli* and *haemolytic E. coli* in the weaned pigs following EHEC challenge. Moreover, COS inclusion in feed alleviated inflammatory responses and disruption of the intestinal epithelium in the EHEC-challenged pigs.

## Data Availability

Data can be made available upon reasonable request to the authors.

## Authors’ contributions

Jiahao Liu: Investigation, Writing-Original Draft, Validation, Formal analysis. Simona De Blasio, Liam Good, Ludovic Pelligand and jun he: Conceptualization, Investigation, Writing-Reviewing and Editing, Visualization, Data Curation, Project administration. Kunhong Xie, Xiang Li, Yuheng Luo, Hui Yan, Xiangbing Mao and Quyuan Wang: Resources, Supervision.

## Conflict of Interest Statemen

The authors declare that there are no conflicts of interest

